# Immune Epitopes of SARS-CoV-2 Spike Protein and Considerations for Universal Vaccine Development

**DOI:** 10.1101/2023.10.26.564184

**Authors:** Nicholas Magazine, Tianyi Zhang, Anang D. Bungwon, Michael C. McGee, Yingying Wu, Gianluca Veggiani, Weishan Huang

**Author notes:** These authors contributed equally to this work (Nicholas Magazine, Tianyi Zhang). Correspondence: Address: 1909 Skip Bertman Drive, Baton Rouge, LA 70803, USA.

## Abstract

Despite the success of global vaccination programs in slowing the spread of COVID-19, these efforts have been hindered by the emergence of new SARS-CoV-2 strains capable of evading prior immunity. The mutation and evolution of SARS-CoV-2 have created a demand for persistent efforts in vaccine development. SARS-CoV-2 Spike protein has been the primary target for COVID-19 vaccine development, but it is also the hotspot of mutations directly involved in host susceptibility and immune evasion. Our ability to predict emerging mutants and select conserved epitopes is critical for the development of a broadly neutralizing therapy or a universal vaccine. In this article, we review the general paradigm of immune responses to COVID-19 vaccines, highlighting the immunological epitopes of Spike protein that are likely associated with eliciting protective immunity resulting from vaccination. Specifically, we analyze the structural and evolutionary characteristics of the SARS-CoV-2 Spike protein related to immune activation and function via the toll-like receptors (TLRs), B cells, and T cells. We aim to provide a comprehensive analysis of immune epitopes of Spike protein, thereby contributing to the development of new strategies for broad neutralization or universal vaccination.

## Introduction

SARS-CoV-2, the etiological pathogen of COVID-19, is currently responsible for over 760 million cases of illness and 6.9 million deaths as of July 2023 ^1^. As a single-stranded positive-sense betacoronavirus, SARS-CoV-2 exploits the host machinery for replication ^2^. Spike protein is a key envelope glycoprotein for SARS-CoV-2 infection. The SARS-CoV-2 Spike protein assembles into trimers that can be broadly split into two subunits, S1 and S2, divided by a furin cleavage site (**Figure 1A & B**). The S1 subunit is primarily responsible for initial angiotensin-converting enzyme 2 (ACE2) interaction and binding, while the S2 subunit enables membrane fusion ^3^. The S1 subunit contains a receptor-binding site (RBD), which directly binds to ACE2, as well as an N-terminal domain (NTD), which is considered primarily structural in Spike function ^4^. In contrast, the S2 domain contains two heptad repeat regions (HR1 and HR2) as well as a fusion peptide domain (FP) ^5^. The FP acts immediately after the RBD-mediated Spike-ACE2 interaction by inserting itself into the host membrane, causing a drastic conformational change that brings the HR1 and HR2 domains into proximity with the cell membrane, leading to membrane fusion and viral internalization ^6^.

**Figure 1:**
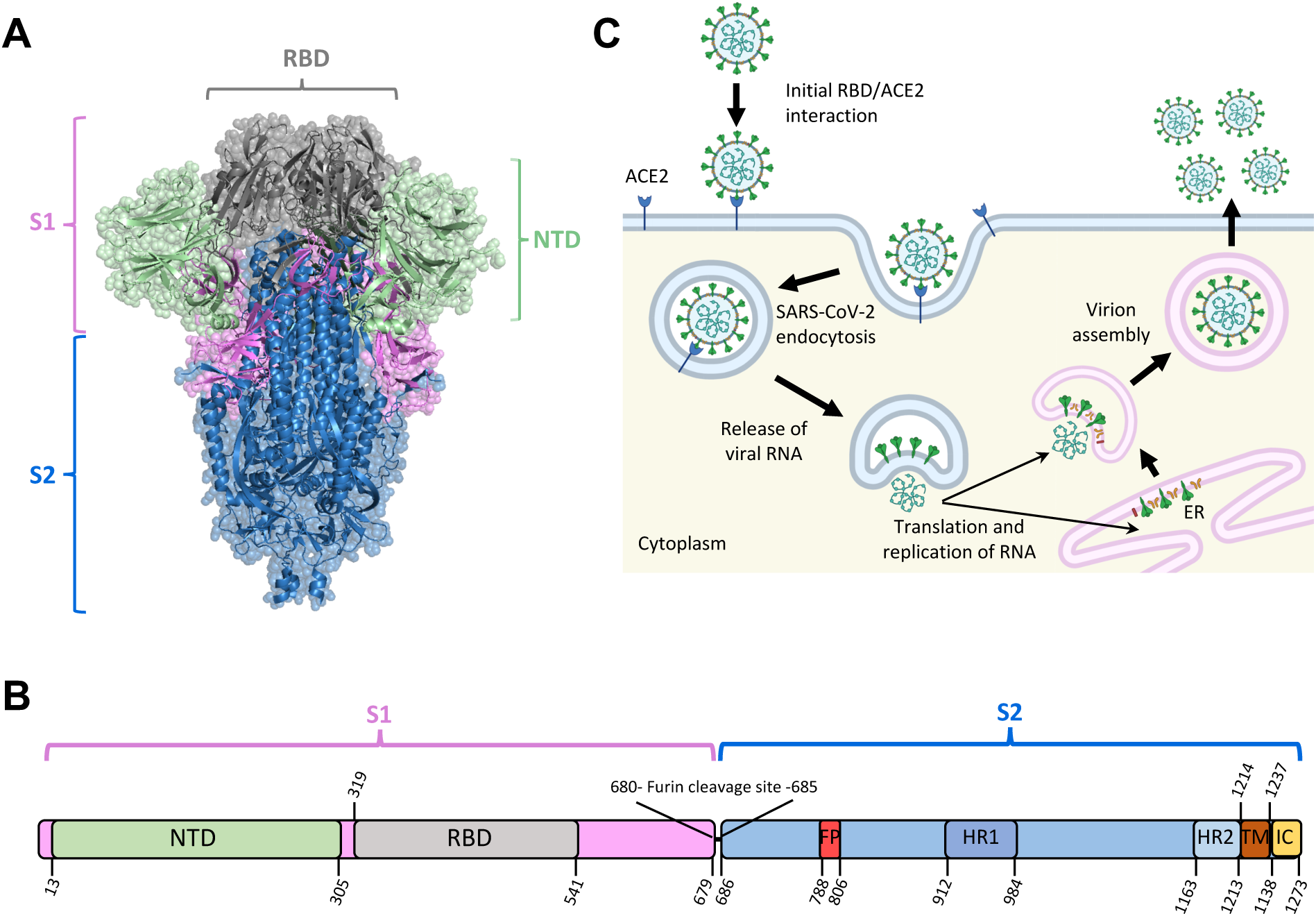
Schematics of Spike structure and SARS-CoV-2 infection. (A) Structure of the SARS-CoV-2 Spike protein (PDB ID 6VXX). (B) Diagram of SARS-CoV-2 Spike (S) gene. NTD: N-terminal domain; RBD: receptor-binding domain; FP: fusion peptide; HR1: heptad repeat 1; HR2: heptad repeat 2; TM: transmembrane region; IC: intracellular domain. (C) The primary mechanism of SARS-CoV-2 cell entry and propagation.

Spike protein recognizes and binds to targeted host receptors, predominantly ACE2 receptor ^3,7^, initiating viral entry into host cells via endocytosis. After entry, the viral RNA escapes the host endosome and exploits the translational machinery of the cell for viral protein production ^8^, followed by assembly and release of the virions from the host cells ^9,10^ (**Figure 1B**). Since it relies on the translational mechanisms of host cells, Spike protein is subject to post-translational modifications (PTMs), which may alter Spike-mediated host-virus interactions and host immune responses. Particularly, glycosylation, the addition of glycan molecules to a protein, has been demonstrated to affect the interaction of a variety of biomolecules with Spike protein ^11^.

Since the identification of its role in the initiation of SARS-CoV-2 infection, Spike protein has been of interest to the scientific community as a primary target for vaccine design and therapeutics ^5,12–14^. The Spike-based vaccine-elicited host immune cell responses can be broadly divided into three arms: innate immune response, in part as a result of toll-like receptor (TLRs) recognition; T cell response; and B cell response ^15–20^. In brief, innate receptor ligands such as TLR ligands in the vaccine can activate innate immune cells, particularly antigen-presenting cells (APCs), which further prime the activation of CD4^+^ and CD8^+^ T cells. Among the activated CD4^+^ T cells, follicular helper T (Tfh) cells contribute to B cell activation, plasma cell maturation, and antibody production, while type 1 helper T (Th1) cells help potentiate APC function, CD8^+^ T cell activation, and cytotoxicity. The vaccine-elicited antibodies provide protection through the neutralization of viruses, while the cytotoxic T lymphocytes (CTLs) eliminate infected host cells to promote clearance of infections (**Figure 2**). The immune subset specialization is heavily influenced by the vaccine adjuvants, while the specificity of the vaccine-elicited antibodies and CTLs is largely defined by the antigenic immune epitopes of the vaccine ^21,22^.

**Figure 2:**
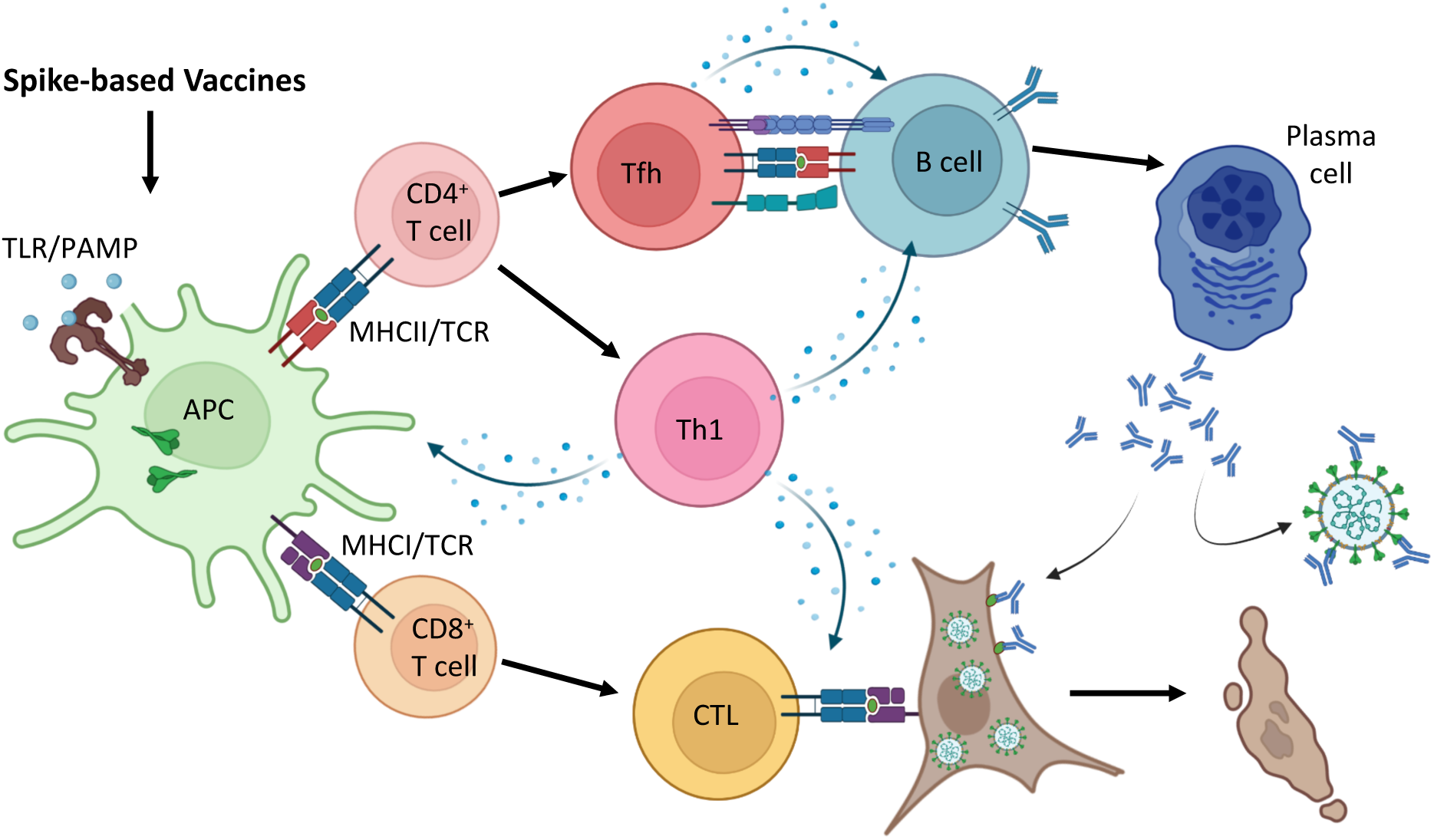
A general overview of host immune responses to Spike-based vaccines. Upon host exposure to Spike-based vaccines, antigen-presenting cells (APCs) sample, process, and present the antigens to CD4^+^ and CD8^+^ T cells via MHCII and MHCI respectively. Activated CD4^+^ T cells can differentiate into follicular helper T (Tfh) and type 1 T helper (Th1) cells which further mediate B cell activation and plasma cell maturation. Th1 cells also aid in priming APC activation and maturation to enhance antigen presentation. Activated CD8^+^ T cells also receive assistance from Th1 cells to potentiate development into cytotoxic T lymphocytes (CTLs). While plasma cell-produced antibodies can neutralize viruses and bind to infected cells to mediate antibody-associated cell death, CTLs function through direct contact to eliminate infected cells. Therefore, Spike-based vaccine-induced antibodies and T cells provide protective immunity against SARS-CoV-2 infections.

Host-virus interactions mediated by Spike protein appear to be the primary driver of SARS-CoV-2 evolution ^23,24^. In the three-year period following the initial outbreak of COVID-19, most circulating SARS-CoV-2 strains have accumulated upwards of 40 mutations in their Spike proteins. Given that most vaccinations administered to the global population specifically target Spike proteins, this may in part explain the sharp decrease in vaccine efficacy over a relatively short period of time ^25,26^. With cases of breakthrough SARS-CoV-2 infections in vaccinated populations increasing recently ^27^, it has become apparent that the development of universal vaccines that are resilient against emerging SARS-CoV-2 variants will be required moving forward.

A universal vaccine is largely defined as one that is safe, long-lasting, and effective against multiple variants ^28^. A detailed understanding of the underlying factors of host immune responses to SARS-CoV-2 is critical for the development of a universal COVID-19 vaccine. Here, we explore the characteristics of immunogenic epitopes on Spike protein, aiming to identify regions of the SARS-CoV-2 Spike protein that are of particular interest in the context of universal vaccine development. Specifically, we explore the interactions between Spike immunogenic motifs and immune receptors on APCs, T cells, and B cells and discuss the efficacy and sustainability of specific T and B cell immune epitopes for universal vaccine development.

### Interactions of SARS-CoV-2 Spike protein with Toll-like receptors

TLRs direct the activation and localization of cells of the innate immune system, ultimately governing the initial host response to both infection and vaccination. Following SARS-CoV-2 infection or the administration of therapeutics that directly or indirectly introduce SARS-CoV-2 Spike protein into a host system, an innate immune response is initiated, in part, via TLR activity ^18,29^. Although this reaction is integral to the host immune response, multiple studies have shown excessive TLR activation to be a driver of uncontrolled inflammation and ultimately tissue injury upon host exposure to SARS-CoV-2 ^30,31^. In the context of SARS-CoV-2 therapies, cell-surface TLRs that are capable of recognizing glycolipids, proteins, and peptides (such as those in SARS-CoV-2 Spike) are of particular importance. The involvement of TLRs, including TLR1, TLR4, and TLR6, in COVID-19 pathogenesis has been demonstrated ^30–33^. TLR2 primarily recognizes lipoproteins, and its role in SARS-CoV-2 Spike interaction has been suggested to be minor and partially dependent on the formation of heterodimers with both TLR1 and TLR6 ^34^. Patients with severe COVID-19 have been demonstrated to have high serum levels of cytokines downstream of the MyD88 signal cascade ^31,35^. Additionally, vaccines with supplemental adjuvants targeting TLR1 and TLR4 have been demonstrated to provide a higher level of protection as compared to controls ^36–38^.

Structure-based predictions have suggested that multiple Spike residues likely interact with TLR1, TLR4, and TLR6 (**Figure. 3A-C**) ^39–46^. Tthe sites predicted to be involved in direct interaction with TLRs are predominantly located in the Spike S1 subunit flanking the RBD, including the NTD and regions to the C terminus of RBD on the S1. Note that there is no preference toward hydrophobic or hydrophilic interaction (**Figure 3D**). All residues interacting with TLR1 and TLR6 can be found within the Spike S1 subunit, with a particular density within regions 30-117 and 204-306. Notably, the Shannon entropy (SE), which represents the degree of genetic variation between all SARS-CoV-2 variants of all residues within this region, is very low, with no single residue exceeding 0.05, even when exclusively considering the Omicron BA.5 lineage (**Figure 3D**). Low genetic entropy indicates a low rate of genetic variability within this region, suggesting that TLR-interacting residues are relatively conserved. Therefore, it is likely that vaccine-elicited immune responses to these regions may induce cross-reactive and long-lasting protection even in the occurrence of new variants.

**Figure 3:**
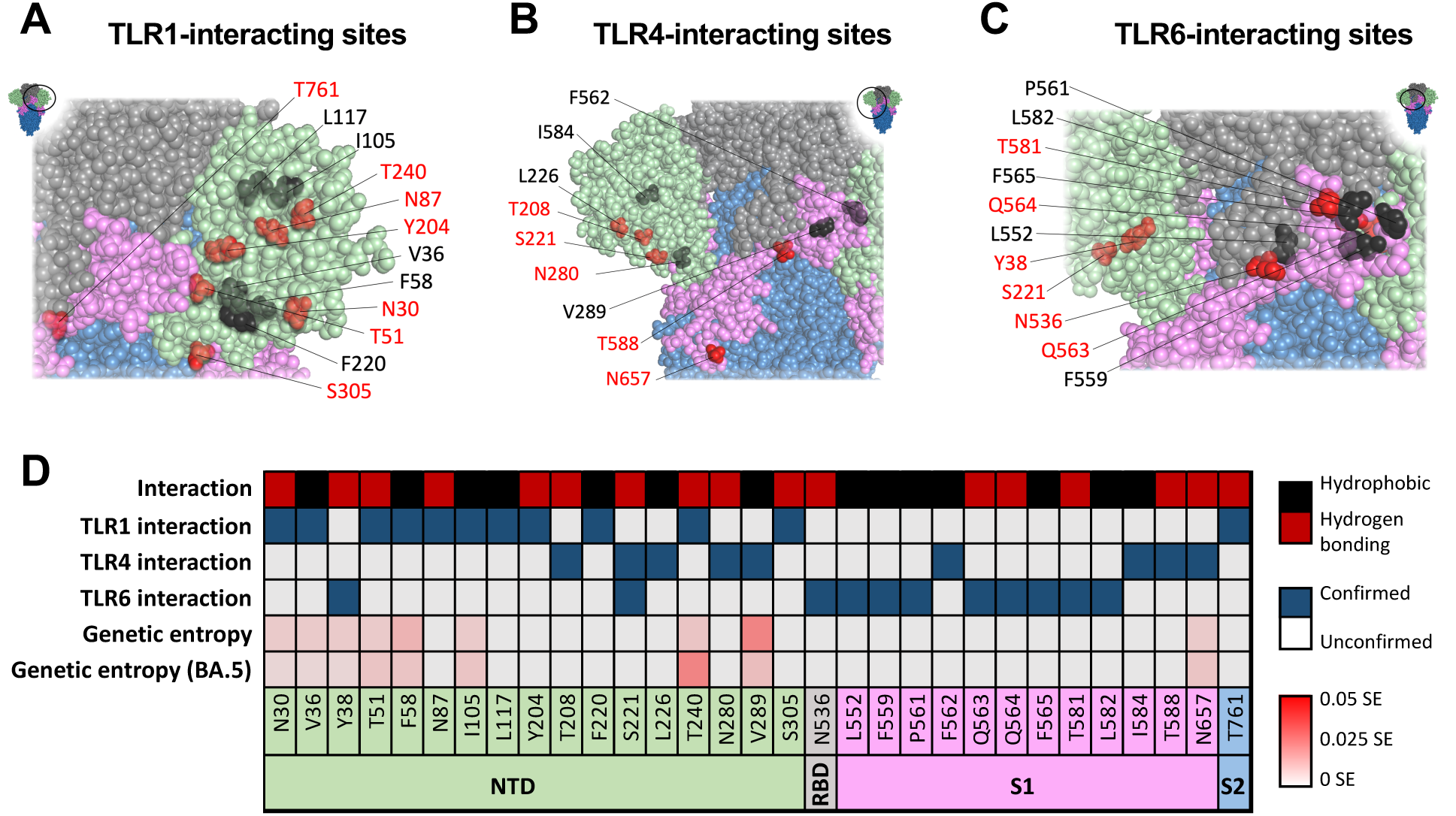
Toll-like receptor interacting epitopes in SARS-CoV-2 Spike. Sites on SARS-CoV-2 Spike protein interacting with TLR1 (A), TLR4 (B), and TLR6 (C). Inset shows full-length Spike protein with a circle indicating the relative position of zoomed in area. Sites interacting via hydrophobic interaction or hydrogen bonding are displayed in black or red respectively. (D) List of the TLR-interacting sites, interaction type, TLR interaction, genetic entropy of all reposited sequences, and genetic entropy relative to Omicron strain BA.5. Blue boxes indicate experimentally confirmed TLR-interacting sites. Sites with Shannon entropy (SE) exceeding 0.05 bear substantial variation among the described SARS-CoV-2 strains.

The described regions are expected to interact with TLRs and help induce an enhanced initial response of APCs that is conducive to higher vaccine efficacy. Although this enhanced initial response may often be beneficial in eliciting host immunity against viruses, hyperinflammation has repeatedly been associated with adverse reactions to vaccination as well as severe COVID- 19 ^47–49^. Therefore, in addition to their potential benefits, these TLR-interacting motifs should also be considered for potential vaccine-associated adverse effects. The inclusion of TLR agonists as adjuvants has been a popular strategy in vaccine development, with numerous demonstrated successes ^50,51^. It is, however, critical to understand the agonistic properties of endogenous TLRs in vaccine antigens to guide the selection of additional adjuvants for emerging diseases. While the inclusion of TLR agonists is likely beneficial to vaccine efficacy for most populations, they pose a potential danger of provoking vaccine-associated acute inflammation mediated by an excessive innate immune response.

### T cell epitopes of SARS-CoV-2 Spike protein

Being a primary driver of the adaptive immune response, T cells modulate humoral and cellular immunity to SARS-CoV-2 infection and play a critical role in the maintenance of long-term SARS-CoV-2 immunity ^17,52,53^. Studies have demonstrated the persistence of T cell-mediated immune responses between SARS-CoV-2 variants of concern (VOCs) ^54–56^, including the Omicron BA.5 strain ^57^. This cross-reactive T cell immunity may be the primary driver of resistance to these newer strains conferred by early COVID-19 vaccines, suggesting a critical role of T cell immune epitopes for universal vaccine development.

As T cells primarily act to limit viral replication by mediating APC activation, humoral immune response, and destruction of infected cells (**Figure 2**), they likely do not prevent SARS-CoV-2 infection. Although the prevention of disease is the ideal outcome of vaccination, the prevention of severe clinical manifestations and the further spread of disease is also a desirable result ^58^. This assertion is bolstered by the greatly decreased incidence of hospitalization for vaccinated patients upon clinical presentation of COVID-19 following Omicron infection, despite having received vaccination for non-Omicron SARS-CoV-2 lineages ^59–61^.

T cells primarily recognize short, linear peptides ranging from 8 – 15 amino acids in length presented on human leukocyte antigen (HLA) molecules ^62,63^. There are two major HLA groups, class I and class II, which present antigens predominantly to the T cell receptors (TCRs) of CD8^+^ and CD4^+^ T cells, respectively ^64,65^. Class II HLA complexes are typically found only on APCs and present peptides from exogenous sources, while class I HLA complexes are expressed ubiquitously and recognize peptides from endogenous antigens ^65^. The linear nature of T cell immune epitopes decreases the complexity of vaccine development targeting T cell receptors, as variation in protein/peptide structure is of minimal concern. However, TCR recognizes peptide antigens only if they are presented by the HLA. HLA genes are expressed in a polymorphic manner, and various populations may be deficient in a specific HLA group ^66,67^. It is speculated that the outcome of TCR-stimulating vaccines is heterogenous, but a potential cross-reactive long-term effect may be achieved in some individuals. A potential strategy to cope with the HLA genetic variance is to identify peptide sequence combinations expressed by the majority of the global population (with some combinations capable of exceeding 99% global population coverage) ^68–71^.

While the binding of a peptide to HLA complex is necessary for the initiation of a T cell response, this is not sufficient. Bioinformatic analysis of T cell epitopes (compiled at the Immune Epitope Database, IEDB) shows a relatively narrow epitope repertoire for T cells as compared to HLA complexes ^72,73^. As of July 2023, a query in IEDB shows a pervasive subset of T cell epitopes (having occurred in ten or more individuals and taking place in over 80% of individuals tested for these epitopes), as demonstrated by the proportion of their incidence among individuals who have been tested for them (**Figure 4A&B**). Interestingly, 9 of these 21 peptides show high average levels of genetic entropy (with a Shannon entropy exceeding 0.05), suggesting that T cell interaction could contribute to SARS-CoV-2 evolution (**Figure 4A**). These 9 epitopes all fall within the NTD and RBD. Structurally both RBD and NTD are critical in SARS-CoV-2 interactions with host receptors. Mutations in S1 including the NTD and RBD are abundant ^23,24^, likely due to their active involvement in host-virus interaction that cause selection pressure imposed host receptors (notably ACE2) and antibodies. Conversely, T cell epitopes in the Spike S2 subunit (residues 751- 767, 936-952, 1011-1028, and 1171-1179) exhibit comparatively lower rates of genetic variability, likely because S2 domain is less engaged in direct interactions between the virus and host receptor/antibodies. Given the relatively low genetic variability, TCR epitopes falling within the S2 region may be target candidates of cross-reactive T cell memory responses for universal vaccine development.

**Figure 4:**
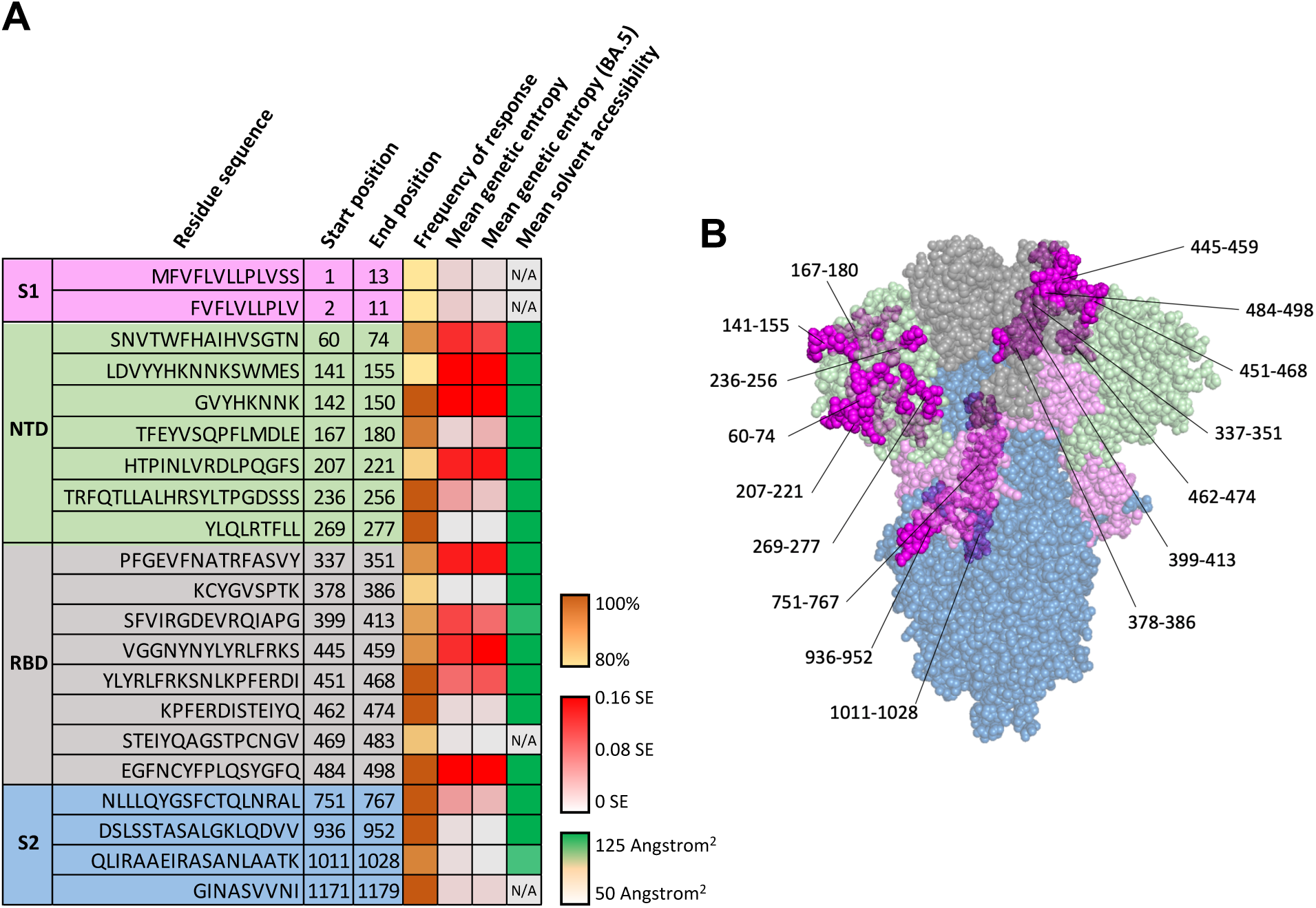
T cell receptor immune epitopes in SARS-CoV-2 Spike. (A) List of T cell receptor interacting epitopes, starting/ending positions, frequency of response as seen in experimental assays (T cell activation or peptide/tetramer binding), mean residue genetic entropy of all reposited sequences, mean residue genetic entropy relative to Omicron strain BA.5, and mean residue solvent accessibility. The frequency of response represents the proportion of positive assays as compared to the total number of assays for a given epitope (higher being more common) with data obtained from IEDB.org. Genetic entropy was calculated based on the Shannon entropy (SE) of GISAID-reposited SARS-CoV-2 sequences and is reported as the mean calculated residue entropy through the given epitope. Solvent accessibility is reported as mean square-angstroms of solvent accessibility for residues through each epitope. (B) Sites on the SARS-CoV-2 Spike reported to have T cell receptor interaction. Surface regions are displayed in magenta; some residues are not on the surface and are occluded by the colored domains, becoming purple or violet; some residues cannot be shown due to their inaccessibility on the structure resolved.

### B cell epitopes of SARS-CoV-2 Spike protein

B cells play a critical role in the adaptive immune response arising from SARS-CoV-2 infection ^15,74,75^. B cells have functions including cytokine production, the secretion of neutralizing antibodies, and antigen presentation. In the context of COVID-19, the primary known protective role of B cells is the secretion of neutralizing antibodies to prevent both initial infection and viral propagation ^76^. Neutralizing antibodies act by binding SARS-CoV-2, primarily Spike protein, and preventing cellular entry of viral particles by sterically blocking their interaction with the host receptors ^77,78^. Anti-Spike antibodies have been the primary scientific focus of COVID-19 pathology and therapeutic development ^79–81^. Experimental analysis of patient sera samples has been used to classify antibodies that are commonly found in individuals infected with SARS-CoV-2. Bioinformatic analysis of anti-Spike binding antibodies revealed over 4,000 antibodies with distinct antigen binding sites that have been isolated from human sera ^72^. While multiple regions of the Spike protein are capable of eliciting antibody responses, only a small subset of these antibodies appears to be capable of inhibiting SARS-CoV-2 infection when used in neutralization assays. The identification of neutralizing antibody epitopes, especially broadly neutralizing antibody epitopes, will aid the development of future therapeutics and universal vaccines.

To best summarize the SARS-CoV-2 Spike protein epitopes recognized by patients’ antibodies, we queried the IEDB database and further cross-referenced our search for antibodies whose neutralizing activity was validated via *in vitro* neutralization assays. This analysis highlighted the distribution and frequency of serum anti-Spike antibodies having linear epitopes as well as the position of each recognized epitope (**Figure 5**). Notably, a large number of the observed antibodies targeted regions throughout the RBD of SARS-CoV-2 Spike protein (**Figure 5A**). However, this observation may partially result from sampling bias created by higher rates of database deposition for antibodies that are found to have neutralizing activity due to their binding to the RBD. Nonetheless, when we compared the relative propensity of regions to which classified serum antibodies bind SARS-CoV-2 Spike protein and the locations of antibody binding sites that have been experimentally validated to effectively neutralize SARS-CoV-2 (**Figure 5B**), there appear to be only three recurring regions to which patient serum antibodies bind and effectively neutralize SARS-CoV-2, the majority of which occur outside of Spike protein’s RBD (**Figure 5B**). Taking into consideration the genetic entropy and solvent accessibility of these regions, many of the epitopes recognized by patients’ antibodies have high solvent accessibility and experience high rates of mutation within the RBD (**Figure 5B**). This observation closely aligns with the notion that mutations in the RBD of SARS-CoV-2 play a part in immune escape from antibody neutralization and thus viral evolution. Interestingly, the neutralizing epitopes of Spike protein that are located outside the RBD appear to be conserved (low genetic entropy), despite their intermediate to high solvent accessibility. In particular, epitopes in the S2 domain (spanning residues 809-876 and 1140-1166) displayed low genetic entropy, high solvent accessibility, and were commonly recognized by patients’ antibodies resulting in good neutralization (**Figure 5B&C**). Therefore, such epitopes would likely be promising targets for universal vaccine development as patient antibodies targeting these areas would likely be abundant, highly neutralizing, and less likely to lose effectiveness against newly emerging SARS-CoV-2 lineages.

**Figure 5:**
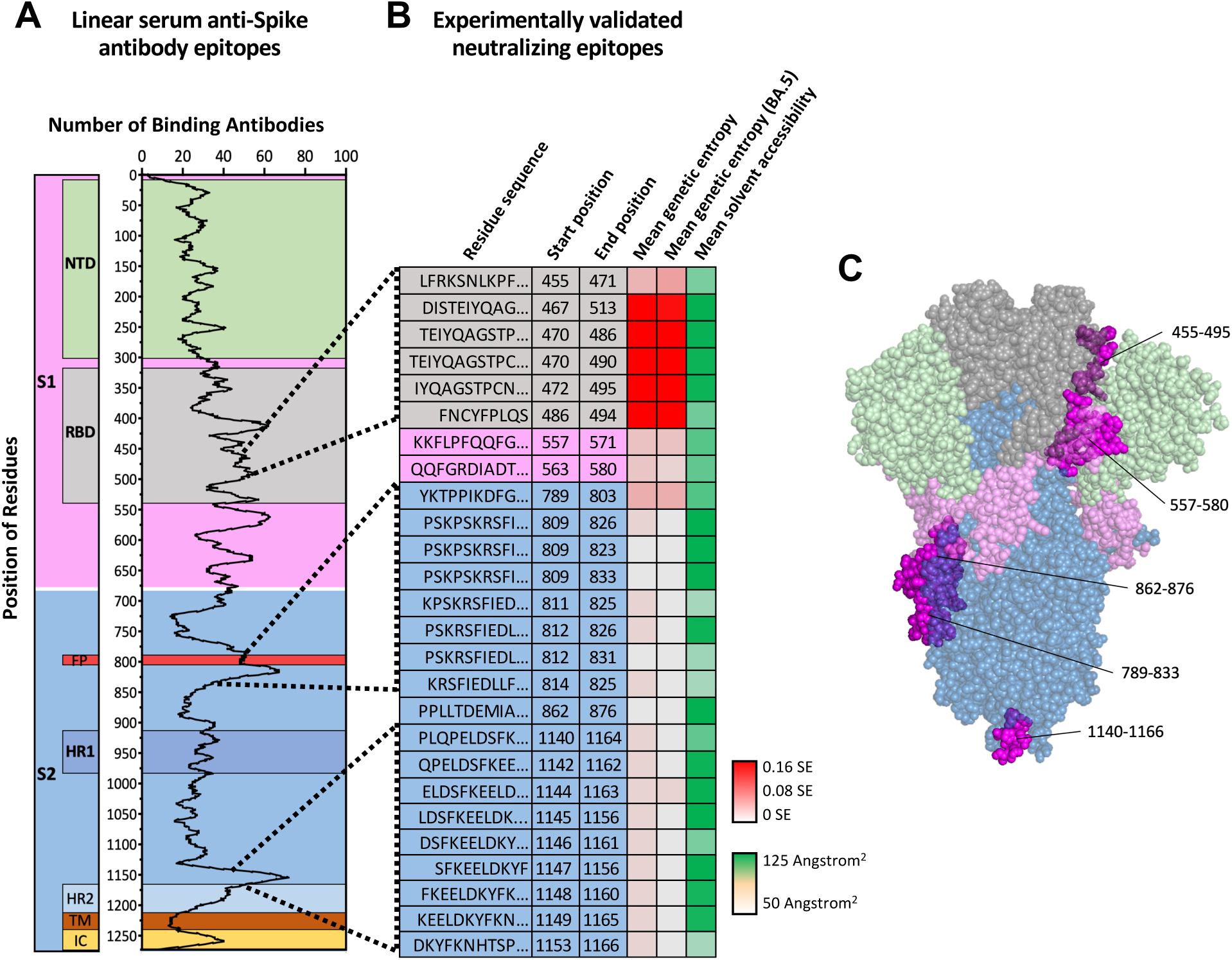
B cell receptor/antibody immune epitopes in SARS-CoV-2 Spike. Comparison of linear epitopes (A) observed as binding targets and (B) experimentally validated as neutralizing antibody epitopes, by antibodies found in database-reposited SARS-CoV-2 patient sera. The first nine amino acid sequences of the epitopes, starting/ending positions, mean genetic entropy of all trains, mean genetic entropy of BA.5, and mean residue solvent accessibility are shown. Genetic entropy was calculated based on the Shannon entropy (SE) of GISAID reposited SARS-CoV-2 sequences. Values are reported as the mean calculated residue entropy through the given epitope. Solvent accessibility is reported as mean square-angstroms of solvent accessibility for residues through each epitope. (C) Sites linear epitopes of neutralizing antibodies on the SARS-CoV-2 Spike protein. Surface regions are displayed in magenta; some residues are not on the surface and are occluded by the colored domains, becoming purple or violet; some residues cannot be shown due to their inaccessibility on the structure resolved.

### Glycosylation in SARS-CoV-2 Spike immune epitopes

Glycosylation is the most common post-translational modification (PTM) occurring within most organisms. During the process of glycosylation, carbohydrate molecules are covalently attached to the side chains of a number of amino acids ^82,83^. Most commonly, this occurs at either oxygen (in Ser, Thr and Tyr) or nitrogen (in Arg and Asn), resulting in O-glycosylation and N-glycosylation, respectively. In eukaryotes, O-glycosylation occurs primarily in the Golgi apparatus, while N-glycosylation occurs primarily in the endoplasmic reticulum ^84,85^. The reasons for which a protein may undergo glycosylation are multifaceted. The added carbohydrate motifs can function as adhesion molecules via interaction with cellular lectins, which may ultimately drive the transportation and sub-cellular location of the modified proteins ^86,87^. Glycans may also obscure the presentation of surface molecules, which may alter the antigenicity of proteins and peptides presented by various pathogens ^88,89^. Finally, glycosylation can substantially alter the structural presentation of a protein by influencing its local conformation as well as its thermostability ^90,91^. During the production of SARS-CoV-2 proteins in host cells (**Figure 1C**), the host molecular machinery also heavily glycosylates viral proteins, including Spike protein. In total, the SARS-CoV-2 Spike protein has been predicted to contain as many as 27 glycosites (predominantly N-glycosylation), the majority of which have been confirmed by mass spectroscopy (**Figure 6A&B**)^11,92–94^.

**Figure 6:**
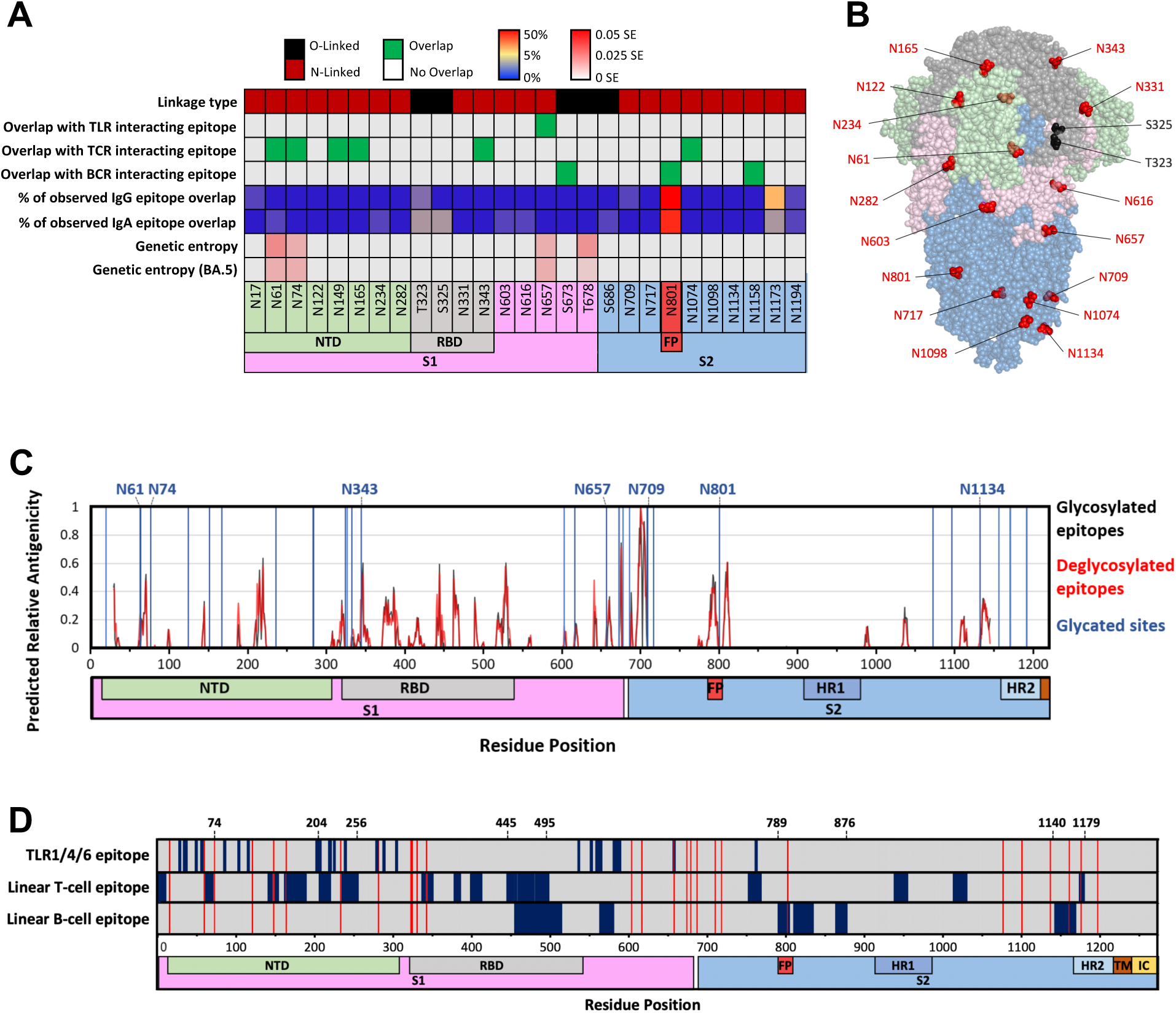
Glycosites and their relations with immune epitopes in SARS-CoV-2 Spike. (A) List of glycosites, linkage type, overlaps with immune epitopes interacting with TLRs, TCR or BCR, proportion of IgG epitope overlap from patient serum, proportion of IgA overlap from patient serum, and genetic entropy of all reposited sequences or only those of the Omicron BA.5 subset. IgA/IgG overlap is reported as the percentage of observed patient-serum antibodies which overlap with a given site. (B) Glycosylated sites of the SARS-CoV-2 Spike protein. Some residues cannot be shown due to their absence from the resolved structure. O-linked and N-linked sites are displayed in black and red respectively. (C) Plots of predicted relative antigenicity over SARS-CoV-2 Spike protein for both glycosylated and deglycosylated Spike. Glycosites are denoted with vertical blue lines. (D) Immune epitope map of SARS-CoV-2 Spike protein. TLR, T cell, and B cell epitopes are displayed. Glycosites are depicted with vertical red lines.

Due to their size, glycans substantially alter the surface accessibility of the residues to which they are attached. This steric hindrance has major consequences for epitope accessibility and ultimately the manner in which a protein is engaged by the human immune system, especially for antibody binding ^95,96^. A comparison of glycosites found on the SARS-CoV-2 Spike protein within immunologically relevant sites reveals that glycosylation occurs at several sites recognized by the immune system. For example, the N801 glycosite falls within a B cell epitope (along with predicted glycosites S673 and N1158) (**Figure 6A**). N657 might interact with TLR4, though this interaction has only been demonstrated computationally (**Figure 3D**). Note that the majority of the glycosites (23 of 27) in Spike protein are highly conserved (see genetic entropy, **Figure 6A**). This conservation may be a result of these glycosites having structural and/or functional roles within the SARS-CoV-2 Spike protein.

Interestingly, N801 has been consistently confirmed as glycosylated ^92^ and demonstrated as a potent epitope for antibody recognition, including IgG and IgA ^72^ (**Figure 6A**). Notably, this glycosite is located within the fusion peptide (FP) (**Figure 6A**) and is also proximal to a region that has been frequently targeted by neutralizing antibodies in the S2 domain (**Figure 5A&B**). An important observation is that the immune epitope containing N801 is well conserved (low genetic entropy) in all VOCs, which is likely associated with its location within the FP region of the S2 domain. The FP is required for virus-cell membrane fusion during viral entry and functionally conserved in coronaviruses ^97^, making it a promising candidate epitope for broad neutralization and universal vaccine design.

In addition to residue N801, several other locations should be noted. Glycosylation sites N61, N74, N149, N165, N343, and N1074 overlap with regions known to engage in TCR interaction, while S673 and N1158 also fall within regions known to elicit neutralization upon antibody binding (**Figure 6A**). Particularly, N1158 falls within a region frequently validated as neutralizing antibody epitope (see **Figure 5B**). It should be noted that not all glycosites are confirmed by mass spectrometry (MS), with O-linked glycosites being particularly erratic in their identification by MS ^93,98–101^.

### Consideration of Spike immune epitopes in universal vaccine development

Taken together, our analyses suggest that regions proximal to N149, N165, N343, N801, and N1158 are immunogenic and relatively conserved. These regions process immune epitopes that can interact with TCRs or immunoglobulins and have low genetic entropy, making them promising antigens for universal vaccine development. Moreover, these regions benefit from low genetic entropy, likely resulting from the structural and/or functional roles of these glycosites. Such a relative conservation would likely reduce the probability of escape from vaccine-induced immunity by future SARS-CoV-2 variants. However, it is possible that this very same glycosylation may also play significant roles in host-virus interactions and the immune responses mounted against these sites. Therefore, it is important to further investigate the effects of glycosylation in peptide-based vaccines targeting these regions, as the presence or absence of a glycan molecule may substantially alter T cell recognition and/or antibody binding. Comparative studies of peptide-based vaccines that include glycosylated versus deglycosylated antigens will provide valuable insights into the impact of glycosylation on vaccine efficacy and immune response.

When it comes to selecting vaccine antigens, it is essential to consider both the immunogenicity of the ‘naked’ antigens as well as their post-translational modifications, as they significantly influence the function and interaction of proteins with the immune system. An example of such steric alterations can be observed in B cell epitopes of the SARS-CoV-2 Spike protein. Linear epitope prediction of the SARS-CoV-2 Spike sequence reveals a wide array of potential binding targets. Nonetheless, it is crucial to assess these targets in the context of their surface accessibility, which contributes to their potential for interaction with immune receptors, especially antibodies. The relative capability of a peptide sequence to bind antibodies can be approximated by multiplying its relative antigenicity by its relative surface accessibility ^102^. To further compare the immunogenicity of the glycosylated and non-glycosylated Spike regions, we computed the relative antigenicity of each spike region using methods adapted from Sikora *et al*^102^. Such computational analyses revealed minimal differences in the predicted immunogenicity of the non-glycosylated versus glycosylated epitopes for eliciting B cell responses (**Figure 6C**).

However, recent studies comparing deglycosylated and fully glycosylated Spike proteins in immunizations of hamsters and mice found that the deglycosylated Spike protein led to enhanced protective immunity ^103^. Furthermore, removal of glycosites in the RBD or especially in the S2 domain resulted in the exposure of more conserved epitopes in both regions, eliciting broadly protective immunity ^104^. These effects may be, in part, contributed by the better accessibility and/or altered stability of the peptide epitopes in the ‘naked’ Spike protein.

## Discussion

A robust understanding of SARS-CoV-2 Spike immune epitopes is critical for the development of universal vaccines and broadly effective therapies that target the Spike protein. Our analysis demonstrates that while an immunological response is observed against nearly every portion of SARS-CoV-2 Spike, only a fraction of these sites elicits a response that is effective in preventing SARS-CoV-2 infection. This observation demonstrates that vaccination using proteins or peptides that target the entirety of Spike protein is likely a less optimal solution when compared to the use of pools of immunogenic epitopes. Such a strategy may not only produce an enhanced immune response to current SARS-CoV-2 variants but will likely provide a stronger and/or more sustainable immunity to emerging SARS-CoV-2 lineages. Our bibliographic and computational analyses indicate that the bulk of a physiologically relevant immune response can likely be elicited by the administration of a subset of Spike protein peptides such as 1-74, 204-256, 445-495, 789-876, and 1140-1179 (**Figure 6D**). Regions 1-74 and 204-256 benefit from having strong interactions with both TLRs and T cells. Regions 445-495 and 1140-1179 contain linear epitopes for neutralizing antibodies and TCR interactions. Both 789-876 and 1140-1179 (**Figure 5B**) benefit from high solvent accessibility and very low relative genetic entropy. The low genetic entropy of these regions suggests that these regions are relatively well-conserved among SARS-CoV-2 variants and unlikely to escape immunorecognition even in the case of novel VOCs. The presence of two glycosylation sites within 1140-1179 makes it less susceptible to mutations. Similarly, region 789-876, housing the glycosite N801 and Fusion Peptide (FP) domain, is likely to remain well-conserved.

SARS-CoV-2 will continue to evolve, and new vaccines to address emerging variants will be needed. This demand will call for the creation of novel vaccines that are effective against a wide range of SARS-CoV-2 variants as opposed to a specific strain. COVID-19 vaccine-elicited antibody responses are critical for protective immunity, but they wane rapidly within a year post-vaccination ^105^. Antigen-specific T cells do not block the initial infection, but induction of T cells specific against conserved domains of SARS-CoV-2 Spike protein demonstrated potential in protecting against SARS-CoV-2 variants that escaped antibody neutralization ^106^. We speculate that administration of peptide pools encompassing regions within the SARS-CoV-2 Spike protein that are conserved and immunogenic, and that can interact with TLRs, TCR and antibodies, will likely produce broad protection from SARS-CoV-2 variants. An important consideration in the design of universal vaccines targeting immunogenic Spike regions would be the potential overlap of these Spike regions with autoantigens. New onsets of autoimmune diseases have been reported post-COVID-19 infection or vaccination ^107,108^, although the underlying mechanisms are unclear. There is an urgent need to further identify and characterize the immune epitopes that overlap with human self-antigens, including self-encoded nucleic acids and proteins, post-translational modifications, and epitopes in the commensal microbiome. Autoantigen-overlapping epitopes should be excluded from future vaccine designs to avoid vaccine-induced adverse effects. Vaccines that contain an enriched pool of conserved and immunogenic epitopes, while avoiding glycosites and autoantigen overlaps will likely not only improve clinical outcomes, but also provide enhanced protection, decrease healthcare costs, improve patient satisfaction, and limit vaccine-associated autoimmune effects.

## Methods

### Genetic entropy analysis

The genetic variability within individual amino acids of SARS-CoV-2 was measured using the Shannon entropy (SE) of codons occurring at the indicated positions. The Shannon entropy value was denoted as ‘genetic entropy’ and was retrieved using the NextStrain (nextstrain.org) COVID analysis tool. Data used for this analysis was obtained from the aligned sequence of SARS-CoV-2 genomes reposited in the GISAID (gisaid.org) as of July 2023. For peptide sequences, the average genetic entropy value of codons throughout the peptide was calculated and reported as ‘mean genetic entropy’. The genetic entropy of all reposited sequences, and the sequences of the BA.5 Omicron lineage, were reported separately.

### Solvent accessibility analysis

Solvent accessibility refers to the solvent-accessible surface area (SASA) of indicated individual amino acids. This value was calculated using the PyMOL by the “Get_area” function, which calculates the surface area in square Angstroms of the selection given with default setting (dot density = 2, solvent radius = 1.4 angstroms, dot_solvent = true). This function was performed for each resolved residue within the closed-state SARS-CoV-2 spike protein (6VXX). For peptides, average solvent accessibility (total solvent accessibility of all resolved residues within the peptide divided by the total number of resolved residues) was denoted as ‘mean solvent accessibility’.

### Experimentally validated epitope determination and frequency of response

Data from the Immune Epitope Database (IEDB.org) was used for the determination of experimentally validated T and B cell epitopes ^109^. T cell epitopes were queried using SARS-CoV- 2 Spike sequence as the epitope source and searching for epitopes that had positive outcomes in T cell assays. These results were further filtered by having human origin, occurring within SARS-CoV-2 and Spike protein, and having linear epitopes. The database was then queried for patient sera samples that had been tested for T cell epitope propensity. The frequency of response for each epitope was calculated by dividing the total number of positive tests by the overall test number. Epitopes demonstrating a frequency of response exceeding 80% and having ten or more reposited tested individuals were reported. Experimentally validated neutralizing B cell epitopes were found by querying the database for SARS-CoV-2 anti-spike antibodies of human origin that have also been tested for SARS-CoV-2 neutralizing capabilities. The linear epitopes of samples meeting these criteria and showing successful neutralization in over 50% of tests were reported. Calculation of the propensity of occurrence for all binding antibodies was performed by querying IEDB for all human-origin antibodies demonstrating SARS-CoV-2 spike binding capacity. Data retrieved July 2023.

### IgG/IgA overlap calculation

Data from the Immune Epitope Database (IEDB.org) was used to calculate the relative propensity for immunoglobulin binding to various Spike regions. A database export containing linear epitopes of human-derived antibodies to SARS-CoV-2 spike between 4-20 amino acids was processed through IgPred Server (iiitd.edu) ^110^ and the resulting IgG/IgA propensity score was reported. Value for each residue has been reported as the mean score for fragments overlapping the target area. Data were retrieved in July 2023.

### B cell epitope antigenicity prediction

B cell epitope antigenicity prediction was performed based on methods used by Sikora *et al* 2021. Accessibility analysis over the SARS-CoV-2 Spike protein (PDB: 6VXX) was performed by rigid-body docking of the Fab fragment of antibody CR3022 ^111^. This analysis was performed on both the native Spike structure as well as the same Spike structure having had its glycans excluded from the model. BepiPred 2.0 webserver was used for the prediction of linear B cell epitopes ^112^. The outputs of these analyses were then multiplied across their respective residues. The maximal output value was then set as 1, and all other values were normalized to the maximum across the entire data set to produce epitope scores for all residues in the predicted epitope regions.

## Acknowledgement

This work was supported in part by grants from the National Institutes of Health (P20 GM130555, R21 AI166898, and R01 AI151139) and a Big Idea Research Grant from the Provost’s Fund for Innovation in Research at Louisiana State University. W.H. received a Research Publication Grant in Engineering, Medicine, and Science from the American Association of University Women.

## Author Contributions

N.M. and T.Z. analyzed and interpreted data, made the figures, and wrote the manuscript; G.V., Y.W., M.C.M., and A.D.B. reviewed and edited the manuscript; W.H. conceived the research, secured funding, made the figures, and wrote the manuscript. All authors approved the final version of the manuscript.

## Data availability

The findings of this study are based on metadata reposited within the GISAID (gisaid.org) and IEDB (iedb.org) databases as of July 2023. The furin-cleaved spike protein of SARS-CoV-2 with one RBD erect structure (PDB, 6VXX) was used to present the 3D structure of Spike protein in this article. The PyMOL Molecular Graphics System, Version 2.0, Schrödinger, LLC, New York, NY, USA) was used for structural analysis and rendering.

